# Information integration for nutritional decision-making in desert locusts

**DOI:** 10.1101/2022.05.16.492099

**Authors:** Yannick Günzel, Felix B. Oberhauser, Einat Couzin-Fuchs

## Abstract

Swarms of the migratory desert locust can extend over several hundred square kilometres, and starvation compels this ancient pest to devour everything in its path. Theory suggests that gregarious behaviour benefits foraging efficiency over a wide range of spatial food distributions. However, despite the importance of identifying the processes by which swarms locate and select feeding sites to predict their progression, the role of social cohesion during foraging remains elusive. We investigated the evidence accumulation and information integration processes that underlie locusts’ nutritional decision-making by employing a Bayesian formalism on high-resolution tracking data from foraging locusts. We tested individual gregarious animals and groups of different sizes in a 2-choice behavioural assay in which food patch qualities were either different or similar. We then predicted the decisions of individual locusts based on personally acquired and socially derived evidence by disentangling the relative contributions of each information class. Our study suggests that locusts balance incongruent evidence but reinforce congruent ones, resulting in more confident assessments when evidence aligns. We provide new insights into the interplay between personal experience and social context in locust foraging decisions which constitute a powerful empirical system to study local individual decisions and their consequent collective dynamics.

## 2 Introduction

Driven by the constant threat of starvation and cannibalism (Bazazi et al. (2008)), bands of juvenile desert locusts (*Schistocerca gregaria*) march through semi-arid environments with fragmented vegetation structures. Usually, locust bands form in arid recession areas when favourable weather conditions allow the development of green vegetation (Cressman (2016); Tucker et al. (1985)). With increasing locust densities at sparse food patches, social cues, like the resulting mechanical stimulation between animals, initiate gregarisation of previously solitarious animals (Lester et al. (2005); Simpson et al. (2001)). This density-dependent phase polyphenism (reviewed by Pener and Simpson (2009)), in which animals transition from avoiding others to approaching them, causes a devastating autocatalytic chain reaction of aggregation that promotes further aggregation and, consequently, population growth (Roessingh and Simpson (1994), Simpson et al. (1999)). Unlike solitarious locusts, which populate recession areas in low densities and forage alone, gregarious locusts march together and - upon depletion of local resources - invade new regions. Nevertheless, despite the vast humanitarian, ecological, and economic impact of this ancient plague (Steedman et al. (1988)), little is known about how locusts utilise different sources of information to guide their devastating foraging campaigns.

The transition from solitary to group foraging also implies significant differences in the type, quality, and quantity of information animals can gather. Generally, information can be obtained through direct interactions with the environment - ‘private information’, or derived from the behaviour of others - ‘social information’ (Dall et al. (2005); Danchin et al. (2004); Davidson and El Hady (2019)). Consequently, solitarious locusts are likely limited to private information, while social information is abundant in swarms. Integrating both information classes, when available, could potentially allow individuals to update their self-obtained ‘beliefs’ with those of surrounding conspecifics. This integration of information may be essential in high population densities where nutritional unpredictability is high and environmental cues for locating potential resources are limited (Despland and Simpson (2005)). Various phase-dependent morphological and physiological changes accompany the transition from a sedentary solitarious foraging strategy with a restricted host-plant range (Despland (2005), Pener and Simpson (2009)) to nutritionally opportunistic group foraging. Notably, a reallocation of growth investment from low-level sensory areas (*e*.*g*., primary olfactory neuropils) to higher processing ones (*e*.*g*., central complex) during the transition from the solitarious to the gregarious phase (Ott and Rogers (2010)) may reflect a shift in environmental complexity, and consequently, the need for increased information processing.

Group-living individuals routinely utilise information produced, intentionally or unintentionally, by others. Foraging is facilitated by sophisticated communication and recruitment signals in eusocial insects, such as the waggle dance in bees (Frisch et al. (1967)) or pheromone trails in ants (Czaczkes et al. (2015)). In other species, the transfer of social information relies on inadvertent cues provided by the behaviour of others (Dall et al. (2005); Danchin et al. (2004)). For example, gregarious insects such as cockroaches and fruit flies collectively select resource patches without active recruitment (Lihoreau et al. (2016, 2010)). In these cases, resource selection relies on retention effects - animals stay longer when more conspecifics are present (Amé et al. (2006); Günzel et al. (2021); Lihoreau et al. (2010); Martín et al. (2021)). The presence of others can indicate both location and quality of food, which offer discrete and graded information, respectively (Dall et al. (2005)). Notably, however, social information can be outdated or noisy in many cases, and here, animals may favour the use of private information (Grueter and Leadbeater (2014)). Moreover, the presence of others at a resource patch may also indicate partial exploitation/depletion, competition, or - as observed by Bazazi et al. (2008) - potential danger through cannibalism. Consequently, the social context can result in preference inversion (Laurent Salazar et al. (2017), Günzel et al. (2021). Optimal foraging decisions, therefore, require individuals within a group to obtain both classes of information and an evaluation of their reliability in conjunction with their acquisition costs (Dunlap et al. (2016)).

A potentially similar optimisation process has been illustrated for multi-modal integration during navigation (Buehlmann et al. (2020); Dacke et al. (2019); Hoinville and Wehner (2018)). The core premise - gaining higher prediction accuracy through merging of information classes weighted by their uncertainty - can also be valid for private and social information sources and has been successfully applied to human decisions (Park et al. (2017)). To our knowledge, empirical animal studies have rarely applied similar frameworks for information integration, potentially due to difficulties quantifying the availability of each information class.

To this end, we investigate how personal experience and availability of social cues impact locust foraging decisions and whether locusts form a consensus, *i*.*e*., collective quorum decisions on one of two possible patches of food. For this, we tracked the behaviour of locust groups varying in size during simple patch choice assays with either equal or different quality. Our study shows that foraging locusts utilize private and social information and optimally update their beliefs by balancing incongruent (opposing) cues and reinforcing congruent ones.

## 3 Methods

### 3.1 Animals

Experiments were conducted with late-instar (4th, 5th larval stage) gregarious *Schistocerca gregaria* (Forskål, 1775) desert locusts. The animals were commercially obtained from a local insect breeder (b.t.b.e. Insektenzucht GmbH, Bad Wörishofen, Germany) at least one week prior to experiments. During this time, animals were kept in the experimental room under crowded conditions with fresh blackberry leaves *ad libitum*. Two nights before experiments, animals were deprived of food and water to increase their foraging motivation.

### 3.2 Experimental setup

The setup consisted of a circular PVC arena (*d* = 90 cm) with a transparent Plexiglas plate covering the top (fig. 1, inspired by Günzel et al. (2021)). We prevented animals from climbing by applying petroleum jelly (Balea Vaseline, dm-drogerie markt GmbH + Co. KG, Karlsruhe, Germany) to the arena wall. Each trial lasted 30 minutes and was recorded with and a single Basler camera (25 fps; acA2040 - 90m; Basler AG, Germany) equipped with an IR long-pass filter, placed above the arena. A custom-made infrared LED (850 nm wavelength) grid and a diffuser plate placed below the arena ensured uniform illumination for the video recordings. An additional light source (Hakutatz LED ring light, 33cm 35W Bi-colour 3200-5600K) illuminated the arena from above. A white paper wall (height: 75 cm) was additionally placed around the arena to prevent animals from visual guidance or disruption throughout the trial. We kept the temperature constant (24 °C - 26 °C) during experiments and cleaned the arena with ethanol and water after each trial.

**Figure 1.**
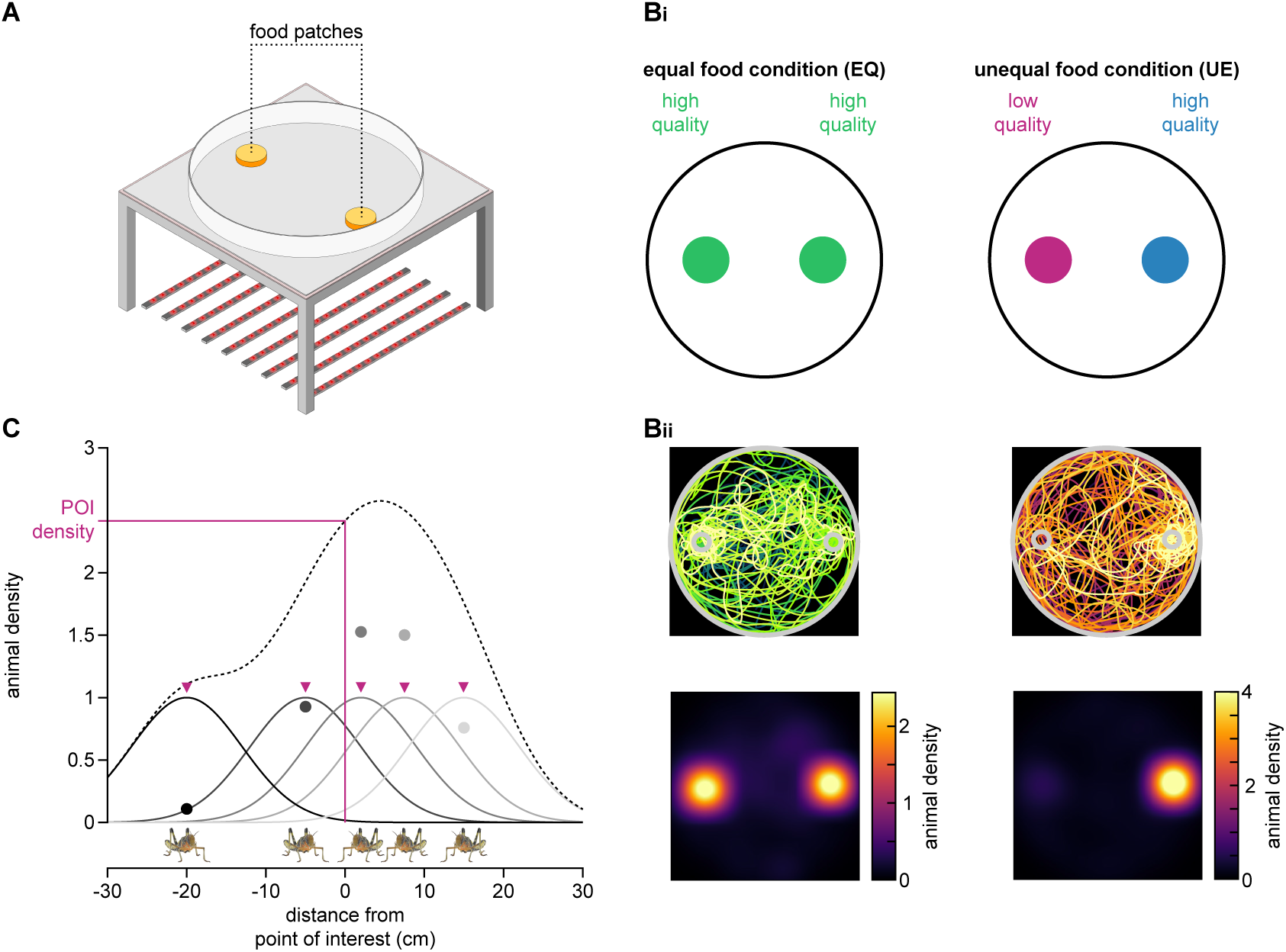
Experimental setup and overview of conducted analysis steps to investigate locust foraging behaviour. (A) Schematic of the circular arena (*d* = 90 cm) used. Two food patches were placed at a randomly selected, opposing position (six possible configurations). (B) Animals were tested under one of two conditions: An equal food condition (*EQ*, left), with two high-quality food patches, or an unequal one (*UE*, right), containing one high quality patch and a low-quality one. Animals were visually tracked using marker-less multi-animal tracking software that reliably maintained individuals’ identities throughout the trial (Bii, first row, example trajectories for trials with *N* = 10 animals, colour codes animals’ IDs). Trajectories were rotated such that food patches were aligned horizontally, with the high-quality patch on the right for *UE*-experiments. Local animal density was estimated based on individual trajectories (Bii, second row, averaged across all frames and animals for the example trials above). (C) Densities were estimated by 2-D Gaussian filtering of individual animal positions within the arena. Shown is a 1-D example with five animals. Solid lines represent local densities induced by each animal. Dots represent local, animal-centric (“perceived”) densities. The dotted line indicates the overall animal density in 1-D space that could be measured at a specific point of interest (POI) as, for instance, the centre of a food patch.

### 3.3 Experimental procedure

Gregarious locusts were tested individually or in groups (*N* = 5, 10, 15, or 30 animals) in one of two experimental conditions: equal trials (“*EQ*-trials”), in which two equal, high quality (“*HQ*-patch”) patches were placed in the arena, and unequal trials (“*UE*-trials”) with one, *HQ*-patch (same quality as in *EQ*-trials) and one with a low nutritional value (“*LQ*-patch”). Food patches were placed opposit each other within the arena in one out of six possible orientations. Under each condition, *EQ* and *UE*, we conducted 20 trials with *N* = 1 animal, ten trials for each *N* = 5, 10, and 15 animals, and four trials with *N* = 30 animals. Thus, 880 animals were tested in total.

### 3.4 Preparation of food patches

For high quality food, rich in water, carbohydrates, and protein, we mixed 500 mL tap water with 38 g gelatin from porcine skin (gel strength 300, Type A, G2500, Sigma-Aldrich, Merck KGaA, Darmstadt, Germany), 25 g honey (Gut & Günstig Bienenhonig flüssig, Edeka Zentrale Stiftung & Co. KG, Hamburg, Germany), and 25 g egg (yolk and albumen stirred beforehand; EDEKA Bio Eier, Edeka Zentrale Stiftung & Co. KG, Hamburg, Germany). The low-quality food consisted of 525 mL tap water and 60 g gelatine, thus providing hydration only to the animals. During the preparation, special care was taken that ingredients mix without caking.

### 3.5 Visual tracking of animals

We obtained trajectories of individual locusts using the multi-animal tracking software TRex (Walter and Couzin (2021)) with a critical focus on maintaining animal identities over time. For video sections with densely clustered animals that were particularly difficult to track, we used a custom-written GUI in MATLAB (R2021a, The MathWorks Inc, Natick, MA, USA) for manual supervision. Obtained trajectories were then smoothed by convolution with a Gaussian kernel (half-width = 2 s, 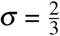s) and rotated to align the patch positions across trials (with the high-quality patch in *UE*-trials always on the right). If an animal stayed for more than 5 s within the range of one body length (4 cm) around the edge of the patch, we considered this interval as a feeding bout. This minimum retention time along with a minimum inter-bout interval of 5 s, accounted for animals just passing by or for noise in tracking, respectively.

### 3.6 Data analysis

We estimated local animal density by a frame-wise application of a 2-D Gaussian smoothing kernel to each animal’s *x/y*-coordinates within the arena. We used a standard deviation that reflected the density-independent interaction range employed by locusts of approx. 7 cm (Buhl et al. (2006)). We estimated the overall animal density at a given point of interest (POI) as the sum of local densities imposed by each animal in the arena (see fig. 1 and mov. 1).

In order to determine the animals’ patch preference, we calculated a preference index based on the local animal densities at both patches at each point in time. Specifically, we divided the difference between patch densities by their sum (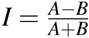; see fig. 2A). Thus, the stronger the preference index differs from zero, the stronger the preference toward a patch. For *UE*-trials, we subtracted the density at the *LQ*-patch from the density at the *HQ*-patch. Thus, positive values reflect a stronger preference for the *HQ*-food and negative values for the *LQ*-food.

**Figure 2.**
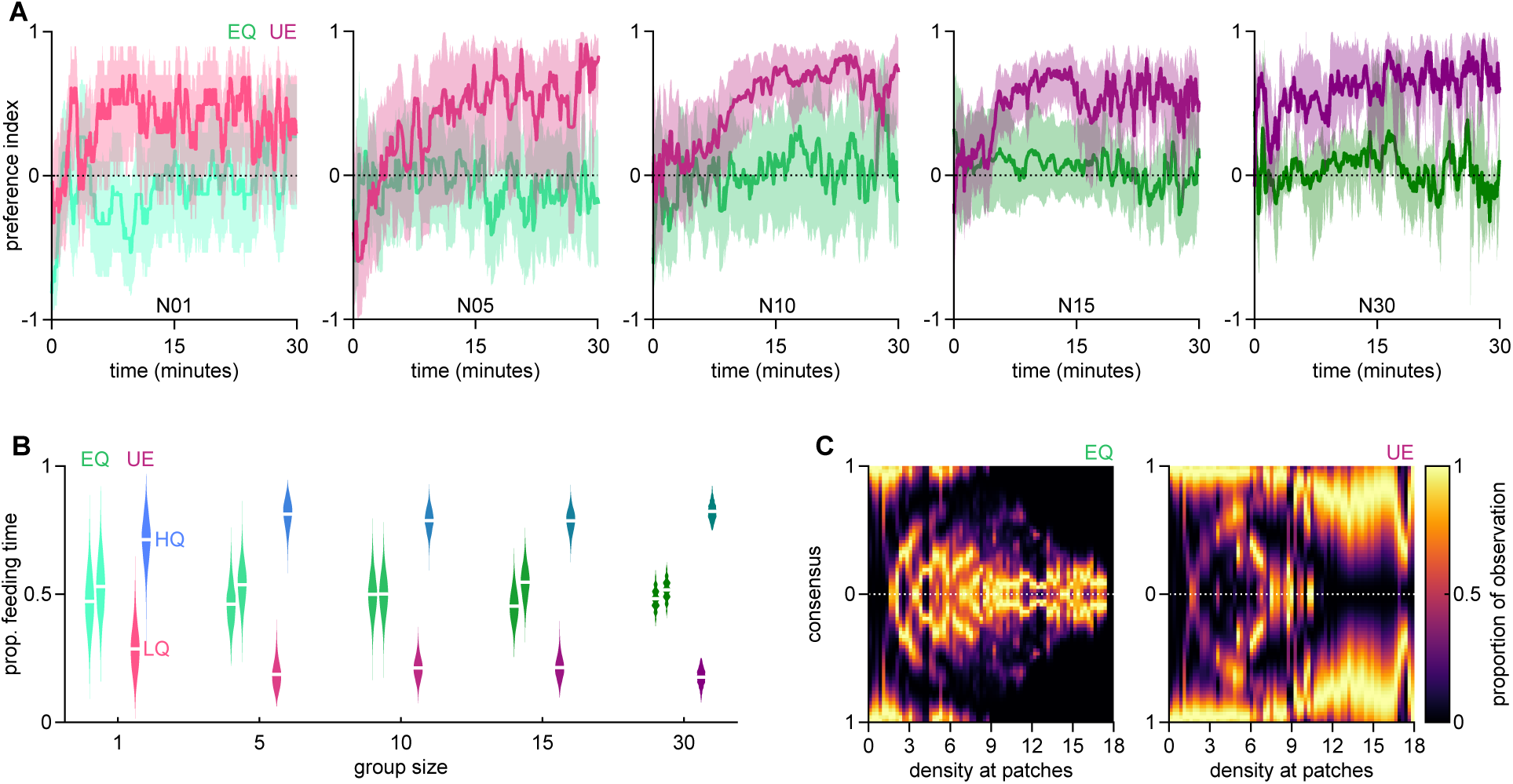
Animals choose food based on quality with group size facilitating their choice. (A) Preference index time courses for the equal (*EQ*; shades of green) and unequal (*UE*; shades of purple) food conditions, separated by group size. The stronger the values differ from zero (dotted line), the stronger the preference towards one patch over the other. For *UE*-trials, positive values indicate a stronger preference for the high-quality option. Solid lines represent grand means across animals and trials. Shaded areas indicate 95% confidence intervals around the mean. (B) Violin plots showing the distributions of the proportion of feeding times at either patch, separated by group size. White lines indicate sample means. (C) Degree of consensus formed by animals as a function of the total animal density at both patches. A value of zero (dotted line) indicates equal density at both patches, while a value of one shows a full consensus - all animals are at a single patch.

To evaluate the distribution of feeding times between the two patches, we calculated the proportion of total time feeding for each patch (see fig. 2B). Bootstrap re-sampling was used to approximate data distributions with *B* = 5’000 samples around the mean.

We were interested in both whether and how animals deal with competition at crowded patches and to what degree competition affects consensus formation. For this, we estimated the degree of consensus as the absolute value of the preference index (see above). For each experimental condition (*EQ* and *UE* trials), we pooled data from all group sizes, binned values according to the amount of prevailing animal density summed across both patches, and normalised by the respective maximum count per bin (see fig. 2C).

To further investigate how the social context affects individual choices, we estimated how joining, staying at, and leaving a patch is associated with animal density at the respective patch (see fig. 3). Joining intervals, *i*.*e*., the interval between two consecutive joining events, were plotted against the average animal density during this interval. Similarly, we plotted stay duration against the average animal density throughout the visit. Here, we binned stay duration and animal density with a bivariate histogram to reveal the respective distributions, as well as to test for a potential interaction between the two. For the latter, we calculated the median stay duration for each density bin, to which we then fitted a linear model. We used the total counts of observations per density bin as weights to account for skewed distributions for the fit. Leaving intervals, *i*.*e*., the interval between a joining and a leaving event at a given patch, were associated with the prevailing animal density at the time of joining.

**Figure 3.**
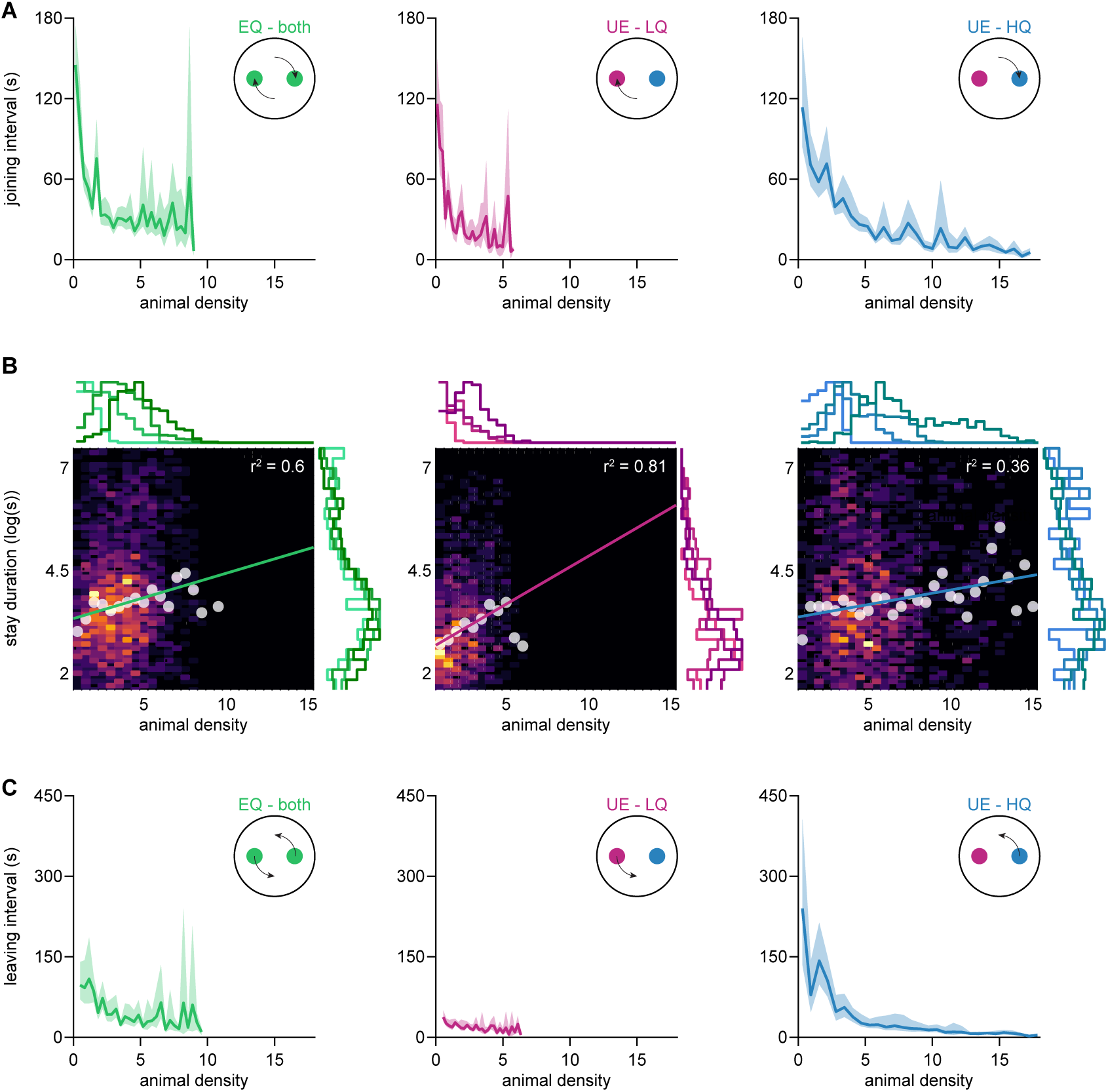
Interactions with food patches were strongly influenced by prevailing animal densities. (A) Interval in between two consecutive joining events as a function of the average prevailing animal density during the interval. Solid lines represent grand means across animals and trials. Shaded areas indicate confidence intervals around the mean. For the equal, high-quality, (*EQ*) condition (left panels), data for both patches were pooled, while for the unequal-quality (*UE*) condition, the low-quality (*LQ*, centre panels) and the high-quality (*HQ*, right panels) were analysed separately. Data were pooled across all group sizes (*N* = 5, 10, 15, 30 animals). (B) Bivariate histograms showing duration of individual visits to a patch as a function of the average prevailing animal density during the stay. Heatmaps illustrate counts per bin. White superimposed dots indicate the median stay duration for each density column, used to fit a linear model to the data (coloured line). Distributions of stay duration and animal density are indicated beside and above the heatmaps, respectively. Individual lines indicate different group sizes ranging from five animals (light colour) to 30 animals (dark colour) per trial. (C) Period between an animal joining a patch and another (or the same) animal leaving it as a function of the prevailing animal density at the joining event. Solid lines represent grand means across animals and trials. Shaded areas indicate 95% confidence intervals around the mean.

### 3.7 Prediction of patch choice

Locust decision-making strategies were analysed by adapting a previously published formulation for Bayesian integration of different classes of information (eqn. 1; Arganda et al. (2012)). The model returns the probability (*P*) of an animal choosing a certain option given both private and social information under the assumptions that (i) animals choose the patch for which they have accumulated more personal evidence, and (ii) animals at a food patch always induce attraction to that patch. The original model included three parameters: the quality of private (asocial) information (*A*), the reliability of social information (*s*), and the relative impact of opposing effects (*k*). However, in our experiments, private information changed over time, depending on the animal’s experience. Tracking all animals with constant identities allowed us to estimate each animal’s amount of accumulated evidence. Thus, we introduced the reliability of private information (*a*) as a new parameter to include the animal’s experience with both food patches:

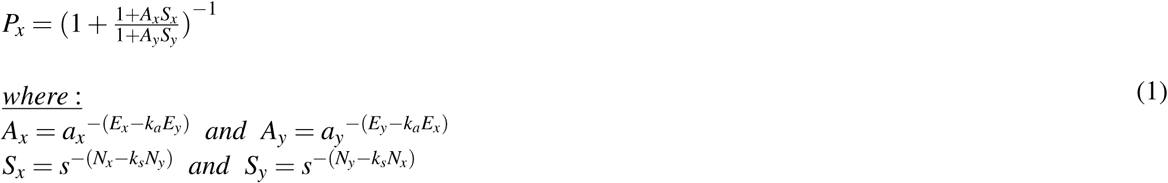

Here, *N*_*x*_ and *N*_*y*_ refer to the accumulated evidence based on social information for patch *x* and *y*, respectively. Likewise, *E*_*x*_ and *E*_*y*_ refer to the accumulated evidence based on private information for patch *x* and *y*, respectively. The model assumes that a patch is a ‘good choice’ and is more likely to be chosen if supporting evidence is substantial. For social information, the more animals of the group were present, the higher the probability of the focal animal joining. Similarly, for private information, the more evidence an animal has sampled during (consecutive) feeding bouts, the higher the probability of it returning. The parameter *k* defines whether animals choose a patch solely based on the relative difference of accumulated evidence (*k* = 1), solely based on the absolute amount of evidence (*k* = 0), or sometimes choose the patch supported by less evidence (0 < *k* < 1), for instance, when crowding is high. We estimated the best-fitting set of parameters for each animal using Bayesian hyperparameter optimisation (using *bayesopt* function in MATLAB) with a lower bound of 1 and an upper bound of 25 for the reliability of information (*a* and *s*). The impact of opposing effects (*k*_*a*_ and *k*_*s*_) was optimised between the bounds of 0 and 1. Distributions of accumulated evidence were root-transformed to account for skewness (SI tab. 2).

Sampling of evidence was modelled as a leaky accumulation process (eqn. 2) with two integrator units per information class, *i*.*e*., one for each patch.

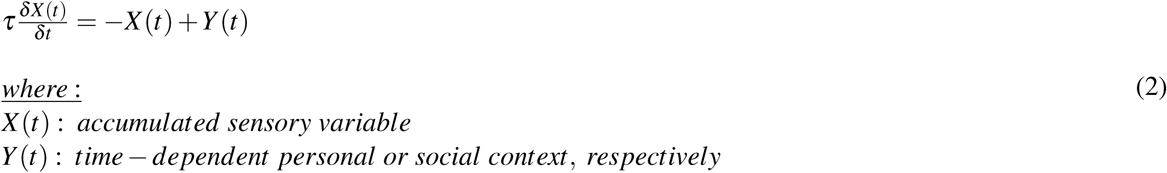

Here, the time-dependent personal context was a step function that equals one for frames in which the animal was feeding and zero otherwise (see fig. 4A, left). The time-dependent social context was the local animal density at a patch with accumulation periods during the focal animal’s resting bouts outside the patch (see fig. 4A, right), based on the finding that resting bouts are associated with reorientation manoeuvres and thus decision-making (Bazazi et al. (2012)). The corresponding time constants *τ*_*personal*_ and *τ*_*social*_ were set to the average integration bout length under the different group sizes (SI tab. 1).

**Figure 4.**
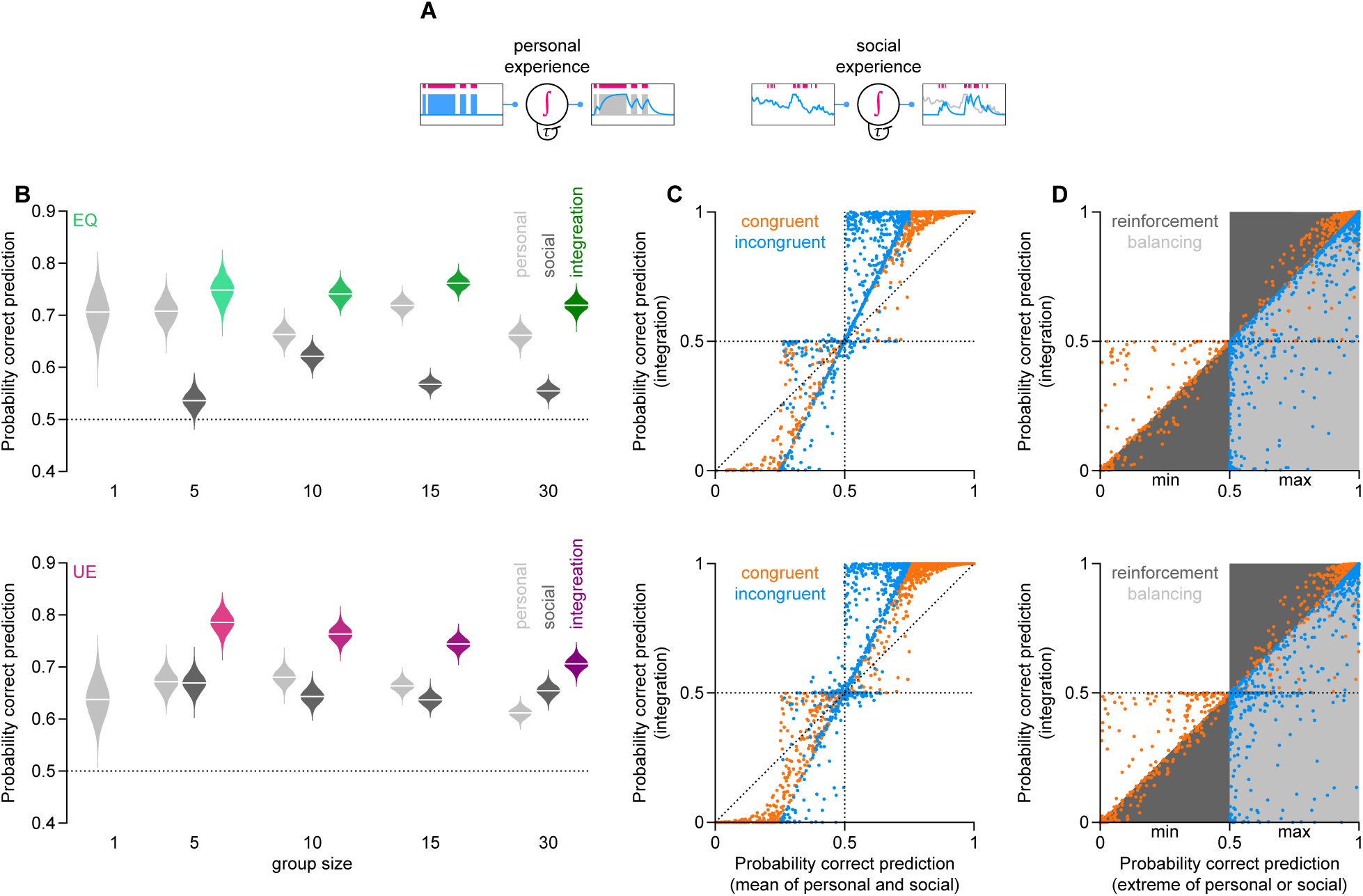
Prediction of patch choice based on Bayesian integration of private and social information. (A) Illustration of the leaky integrator model used for personal and social experience accumulation. Accumulation bouts are indicated in red. We assumed that personally acquired evidence is accumulated during feeding bouts. Socially derived evidence, on the other hand, is assumed to be acquired during reorientation (standing) bouts and based on the prevailing animal densities at the food patches. (B) Violin plots showing the probability distributions of correct patch predictions based on personal information (light grey), social information (dark grey), and the integration of the two classes (colour). Data for the equal food condition (*EQ*) are in the upper row with green violins for the integration and for the unequal food condition (*UE*) below with magenta violins for the predictions based on integration. (C) Scatter plot of the probability of correct predictions from integrating both information classes as a function of the mean between the two single-class (personal or social) predictions. Data points across all group sizes were pooled. Points above 0.5 (dotted horizontal line) reflect correct predictions after information integration. Points above the dotted diagonal line reflect instances where the integration performs better than taking the average of the underlying information classes. Dot colour indicates whether the personal information was congruent (orange; *P*_*private*_ > 0.5 & *P*_*social*_ > 0.5 or *P*_*private*_ < 0.5 & *P*_*social*_ < 0.5) or incongruent (blue; *P*_*private*_ < 0.5 & *P*_*social*_ > 0.5, or *P*_*private*_ > 0.5 & *P*_*social*_ < 0.5) with social information. Data for *EQ* trials are shown in the upper row, with data for *UE* trials below. (D) Scatter plot showing the probability of correct ‘integrated’ predictions as a function of the extreme value (min. or max.) of the single, personal or social, predictions. For the case of congruent information (orange), the smaller value is plotted when both classes mispredicted the patch choice (*P*_*private*_ > 0.5 & *P*_*social*_ > 0.5) while the larger of the two predictions is plotted when both classes predicted correctly. (*P*_*private*_ < 0.5 & *P*_*social*_ < 0.5). In all cases of incongruent information (blue), the larger value is plotted. Shaded areas indicate reinforcement (dark grey) of information, *i*.*e*. integration produces more extreme values when the underlying classes agree, and balancing (light grey) of information when the predictions of the underlying classes conflict. Shown are data points across all group sizes. Points above 0.5 (dotted horizontal line) reflect correct predictions after information integration. Data for *EQ* trials are shown in the upper row, with data for *UE* trials below.

Interested in how locusts make decisions, we dissected the relative contribution of private and social information in the integration process. To do so, we simulated the absence of information by setting the exponents in eqn. 1 to zero. This step either turned the quality of private information *A* or the quality of social information *S* into a value of one, resulting in the absence of private and social information, respectively. Exemplary cases for which this would occur naturally are the absence of private information before the animal’s first feeding bout or ambiguity in social information when other animals are distributed equally among the two options.

### 3.8 Data and code availability

Data were analysed and plotted using custom-written MATLAB scripts. Analysis code and processed trajectories are available at https://github.com/Couzin-Fuchs-Lab/Guenzel-et-al-CollectiveForaging. Custom-written GUI for manually supervised tracking is available at https://github.com/YannickGuenzel/BlobMaster3000. Data are available on request from the authors.

## 4 Results

### 4.1 Condition-dependent formation of consensus

We studied locust foraging dynamics in a simple behavioural assay in which gregarious locusts were tested in different group sizes (*N* = 1, 5, 10, 15, 30 animals) against two food patch options (fig. 1A, mov. 1) of either equal or unequal nutritional quality (fig. 1Bi). We tracked animals while retaining accurate labels of individual identity for 30 minutes to estimate (history-dependent) interactions among conspecifics and with the food patches (fig. 1Bii). Throughout the trials, animals explored the arena with multiple visits to both patches per animal (SI fig. 1A). As locust behaviour is strongly density-dependent, we estimated animal density profiles across the entire arena to characterise each animal’s social experience within, around, and outside the food patches (fig. 1C).

Within a few minutes, animals consistently displayed a persistent preference for the high-quality patch in the nutritionally unequal condition (fig. 2A). We observed this across all group sizes, with increasing coherence as group size increased. The clear preference was also reflected in the proportion of feeding time at the high-quality option (fig. 2B). Note, however, that the proportion of feeding time on the low-quality food was not zero, indicating that animals sampled the two options (cf. SI fig. 1A). Moreover, given how the degree of consensus changed with increasing densities at food patches, a large number of animals interacting with the food appears to have promoted quorum decisions (fig. 2C, for density values above 9). For the nutritionally equal condition, on the other hand, animals did not develop a preference for one patch over the other. Here, an average preference index of zero (fig. 2A), as well as equal feeding times (fig. 2B), could have indicated that animals preferred one patch for one trial but the other in the subsequent trial. However, consulting the consensus maps (fig. 2C) resolves this ambiguity: animals split into two groups already under low densities (large proportion of observations around a consensus value of zero indicate splitting). Taken together, this suggests that desert locusts do not form a quorum decision on a single food patch when an equal alternative is available nearby.

### 4.2 Presence of others dictates individual patch interactions

To better understand the autocatalytic nature of locust swarms, we investigated how the presence of others affects decisions. Tracking individual animals while maintaining their identities throughout the trials allowed us to evaluate how the prevailing animal density affected the probabilities of joining, staying, and leaving a patch. Regardless of the nutritional patch quality, we found that animals joined patches faster (fig. 3A) and stayed longer (fig. 3B) when local density was high. At the same time, however, high density inherits high competition, resulting in shorter leaving intervals (fig. 3C). In other words, a joining animal reduces the time until another (or the same) animal would leave when density is high. Notably, leaving intervals for low-quality patches were short even under low densities, suggesting that leaving events are driven primarily by the low nutritional values rather than competition. This is in line with the generally short bout duration for this patch (SI fig. 1C). Taken together, the density-dependent joining behaviour exhibited in both food conditions suggests that the presence of others serves as an inadvertent social cue, which conveys discrete (location) information but not graded (quality) information. Thus, locusts are required to directly sample (generate private information) to assess nutritional quality. However, can personally acquired evidence update an animal’s belief that has been formed based on a socially derived one - and vice versa?

### 4.3 Locusts integrate private with social information for efficient foraging

Information about the location and quality of food patches forms the basis of foraging decisions. Here, we dissected these decision-making processes by modelling personal and social evidence accumulation. A Bayesian formalism illustrates that locust decisions can be well predicted from the relative difference of accumulated within-class evidence (SI tbl. 2; *k*_*a*_ and *K*_*s*_ approaching a value of one), and by employing an optimal integration of different information classes (private and social information).

For *EQ*-trials, predicting an animal’s choice solely based on private information (fig. 4B; top row, light grey violins) resulted in generally higher probabilities predicting correctly than those solely based on social information (fig. 4B; top row, dark grey violins). The same holds true for *UE*-trials with groups sizes of up to 15 animals. In relatively large groups of 30 animals, social information gives correct predictions with a higher probability than private information. Moreover, the social context in *UE*-trials generally gives more reliable predictions than in *EQ*-trials. This is reflected by a generally higher reliability of social than private information under this condition (SI tbl. 2).

Above all, the integration of both information classes under either nutritional condition (*EQ* or *UE*) exceeds the single cue probabilities of correct predictions (fig. 4B; coloured violins). Personally acquired and socially derived evidence can be congruent, *i*.*e*., both predicting correctly (*P* > 0.5) or incorrectly (*P* < 0.5), or they can be incongruent with one correct and one incorrect prediction (fig. 4C). Yet, the integration of private and social information exceeds the average of the single cue probabilities for most predictions (reflected by the proportion of data points above the diagonal in fig. 4C; *EQ*_*congruent*_ = 78.13%, *EQ*_*incongruent*_ = 81.03%, *UE*_*congruent*_ = 78.41%, *UE*_*incongruent*_ = 75.24%).

When evidence is incongruent there is a conflict regarding the information on which to base a decision. The Bayesian framework adopted here suggests that locusts optimally update their beliefs by balancing conflicting cues (fig. 4D, blue circles). Here, the probability of correct predictions exceeded the chance level in 77.03% (*EQ*-trials) and 74.13% (*UE*-trials) of the patch choice predictions. The same integration rule with congruent evidence (fig. 4D, orange circles) results in reinforcement and reduction of uncertainty, such that the probability of a correct prediction is often more extreme than both single cues. For correct predictions (*P* > 0.5) based on congruent evidence, 73.24% of the *EQ*-trials and 78.89% of the *UE*-trials had a higher probability of being correct than the maximum of the personal and social evidence. This also indicates that incorrect predictions based on congruent cues (fig. 4D, orange circles) could never be rescued (62.08% and 60.32% of congruent incorrect choices for *EQ* and *UE* trials, respectively, had a lower probability of being correct than the minimum of either personal or social evidence). However, since most cases of congruent evidences were correct (less likely that both information classes are wrong) this resulted in more certain correct answers for both nutritional conditions (*P* > 0.5, *EQ* : 55.14%; *UE* : 59.23%). Taken together, modelling locust decision-making processes suggests that animals balance incongruent and reinforce congruent information. This reflects an optimal integration of different classes of information to form stronger opinions when evidence aligns and to consider the respective strengths of the single cues in case of conflicting evidence.

## 5 Discussion

This study provides a framework for studying collective decision-making based on evidence accumulation in different information classes. We show that locusts collectively reach a consensus on the best available food but split into two groups if both options are equal. Our data suggest that these fission-fusion dynamics originate from discrete information on the location of food conveyed by the retention of others. However, by estimating the relative evidence provided by either class of information, private or social, we show that locusts balance incongruent evidence based on their respective strengths. At the same time, they reinforce congruent information to form stronger opinions when evidence aligns.

Reliable sensory-motor decisions rely on stochastic processes to sample information from noisy environments (Davidson and El Hady (2019)), to integrate evidence from different senses (Buehlmann et al. (2020)) according to the internal state (Sareen et al. (2021); Vogt et al. (2021)), and ultimately to generate an appropriate context-dependent motor output (Ache et al. (2019)). Integration can occur at different scales: from the dendritic integration of signals arriving at a single neuron (DasGupta et al. (2014); Groschner et al. (2018)), over small pooling networks of several cells (Borst and Bahde (1988)), up to whole brain regions (Bahl and Engert (2020)). Here, we followed the principles of a drift-diffusion model, a standard model for perceptual decision-making strategies (reviewed by Ratcliff et al. (2016)), by assuming a leaky accumulation of evidence for the location of food (fig. 4). We based the accumulation processes on private and social information, which resemble higher-order constructs of different modalities. Social information in locusts is likely to be based largely on vision, but auditory, haptic, and olfactory cues may also contribute. For simplicity, we consolidated the different aspects of this multi-modal construct into one term, ‘animal density’ (fig. 3). Private information about the quality of a food item, on the other hand, is most likely to be gustatory. Animals were able to remember and discriminate the location of a ‘good’ food patch from that of a ‘bad’ one. This indicates that locusts can accumulate gustatory evidence for several items, store it over seconds, and retrieve the information effectively. Future experiments may elucidate the relative contribution of different modalities and the mechanisms underlying their accumulation processes.

When assessing an event’s likelihood, subjective knowledge and objective facts about such likelihoods are indistinguishable for the observer. Subjective confidence for the occurrence of a given event can be described in terms of Bayesian probabilities, which represent the degree of an observer’s belief (reviewed by Meyniel et al. (2015)). Different classes of information can be combined to increase confidence while making a decision, often according to their inherent uncertainty levels (Cheng et al. (2007)). Information integration processes can explain elevated performance levels in insect navigation and foraging (Buehlmann et al. (2020); Dacke et al. (2019); Hoinville and Wehner (2018); Legge et al. (2014); Wystrach et al. (2015)), human perception (Ernst and Banks (2002)) and collective decision-making (Park et al. (2017)). Here, we show that a Bayesian formalism can also reliably predict foraging decisions by integrating the different information classes (fig. 4).

Predicting locust behaviour is crucial for taking preventive measures against future outbreaks and applying directed control efforts against existing ones. To date, however, forecasts of swarms and marching bands’ locations still fall short of precision (Sword et al. (2010)). Quantitative approaches have been largely insightful for generating testable predictions about the gregarisation process (Collett et al. (1998); Georgiou et al. (2021)), swarm movement (Bernoff et al. (2020); Topaz et al. (2008), and foraging (Lihoreau et al. (2017)). Large-scale empirical studies to corroborate them, however, remain scarce (but note Despland and Simpson (2000a,b) for long-term recordings of gregarisation experiments). Utilising a simple foraging assay, in which the personal and social experiences of all individuals are monitored across time, our study provides an empirical confirmation for the prediction that gregarious behaviour can improve foraging success in locusts (Georgiou et al. (2021)), and shows this behaviour to be dynamically amended by personal experience.

The basic framework for evaluating decision processes based on private and social information could be extended to broader social and spatial scales. The extreme polyphenism of locusts and the fast continuous transition between the gregarious and solitarious phase additionally allows tuning of the valence of social parameters (Pener and Simpson (2009)). With the high-resolution mapping of personal experience, nutritional state, and behaviour, locust foraging offers a powerful empirical system to investigate local individual decisions and their resulting collective dynamics.

## 6 Acknowledgements

The authors thank J. Gübel, Y. Hertenberger, J. Roller, and M. Spagnuolo for help during data collection; as well as A. Land and T. Walter for help tracking animals. This work was completed with the support of the Deutsche Forschungsgemeinschaft (DFG, German Research Foundation) under Germany’s Excellence Strategy – EXC 2117 – 422037984

## 7 Declaration of Interests

The authors declare no competing interests.

---

**Movie 1.**
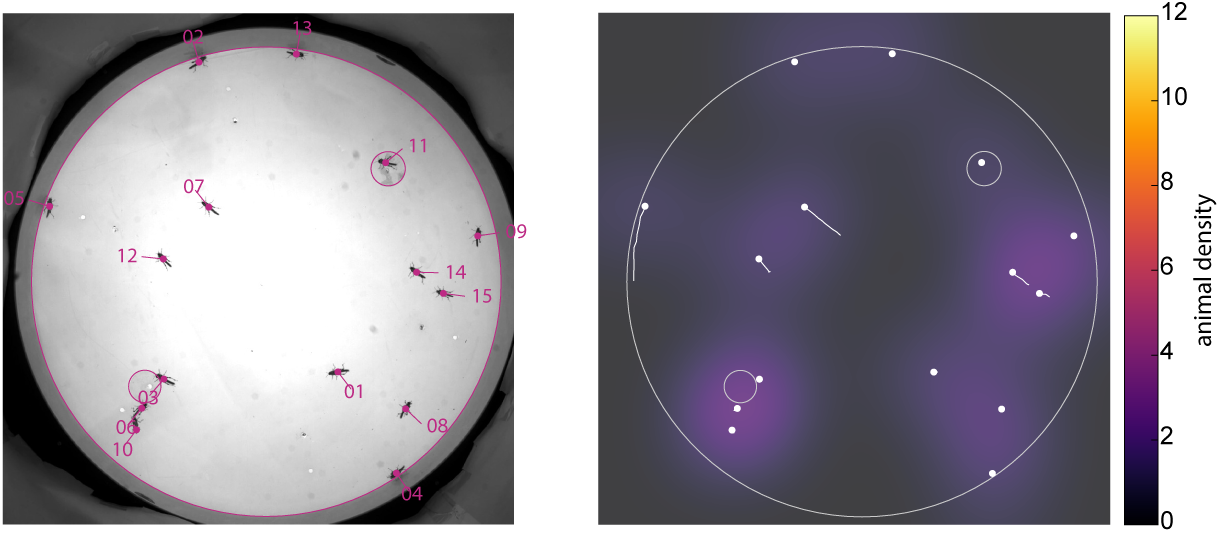
Example foraging behaviour under the unequal food condition. A group of 15 animals sampled both food patches over time but eventually formed a consensus at the high-quality one. The left panel illustrates the accurate tracking of individual identities over time, while the right panel shows the resulting animal density within the arena together with each animal’s centroid (white dots) and trajectory over the past two seconds (white lines). The low-quality patch was placed in the top-right corner. The high-quality patch was placed in the bottom-left corner.

## 8 Supplementary Information

**Supplementary figure 1.**
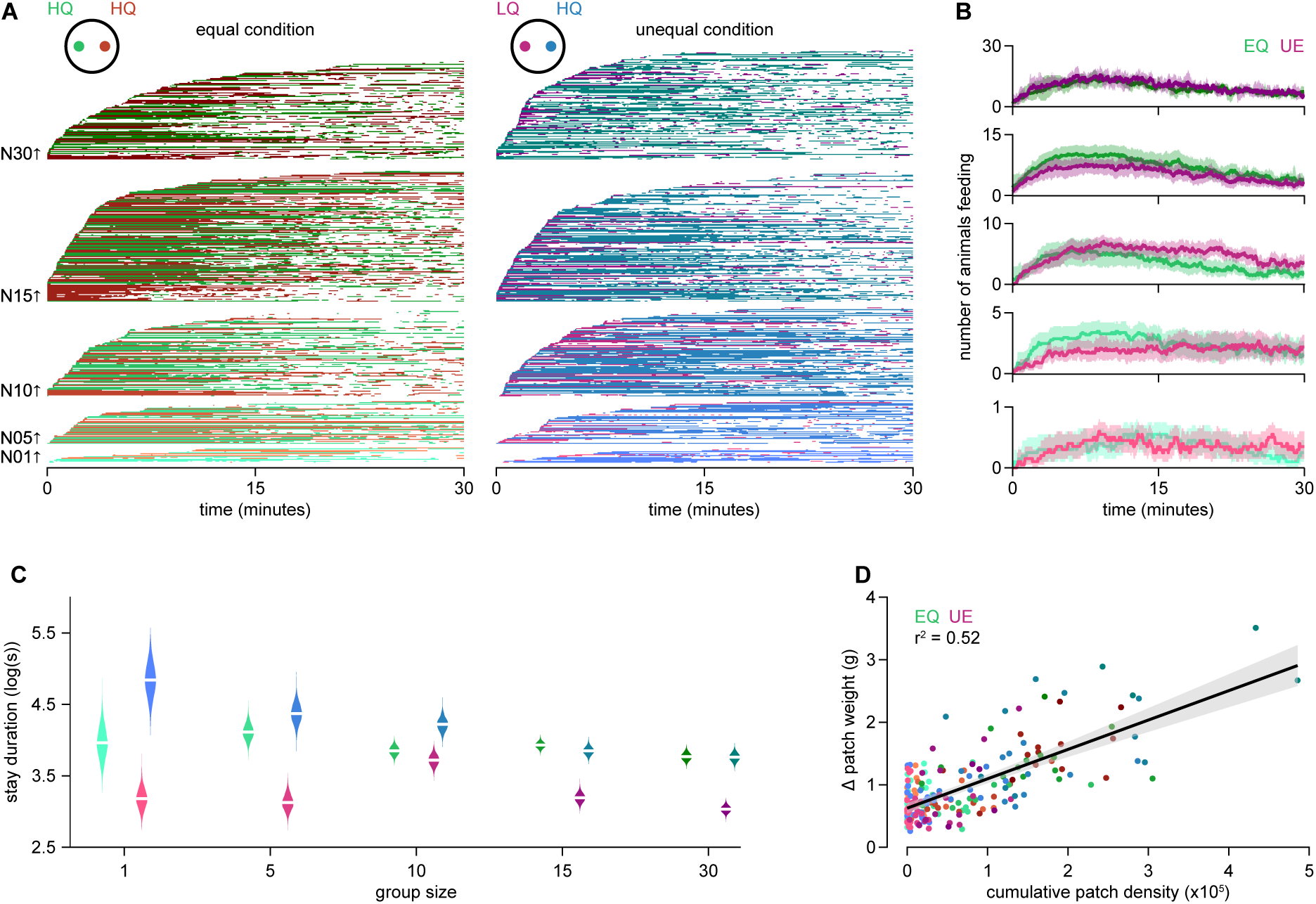
Feeding events. (A) Feeding bouts over time. Each line represents an animal’s repeated visits to either of the patches (colour-coded). Data from each group size (*N*01 to *N*30) were sorted by the occurrence of the first feeding bout. Left: equal food conditions (*EQ*) with two high-quality (*HQ*) food patches. Right: unequal food conditions (*UE*) with one(*HQ*) food patch and one low-quality (*LQ*) food patch. (B) Average number of animals feeding, regardless of patch identity. Shaded areas indicate confidence intervals around the mean. (C) Distributions of stay duration at food patches for each group size. For (*EQ*) trials, data from both patches were pooled (green violins). For (*UE*) trials, magenta violins represent (*LQ*) food patches, whereas blue violins represent (*HQ*) food patches. White lines indicate sample means. (D) Change in patch weight (before-after) of each trial as a function of cumulative patch density over the whole trial. Food patch conditions were colour-coded as in (A). Shaded area indicates the predicted confidence interval around the mean.

**Supplementary table 1.** Time constants [s] for the personal (*τ*_*personal*_) and social (*τ*_*social*_) information class, respectively. Values were estimated for each group size (*N*01 to *N*30) and across both food conditions (*EQ & UE*), based on the average accumulation interval of the respective channel.

|  | N01 | N05 | N10 | N15 | N30 |
| --- | --- | --- | --- | --- | --- |
| $\tau_{personal}$ | 332.10 | 189.83 | 218.85 | 153.20 | 127.51 |
| $\tau_{social}$ | n.a. | 11.19 | 9.07 | 12.42 | 19.25 |

**Supplementary table 2.**
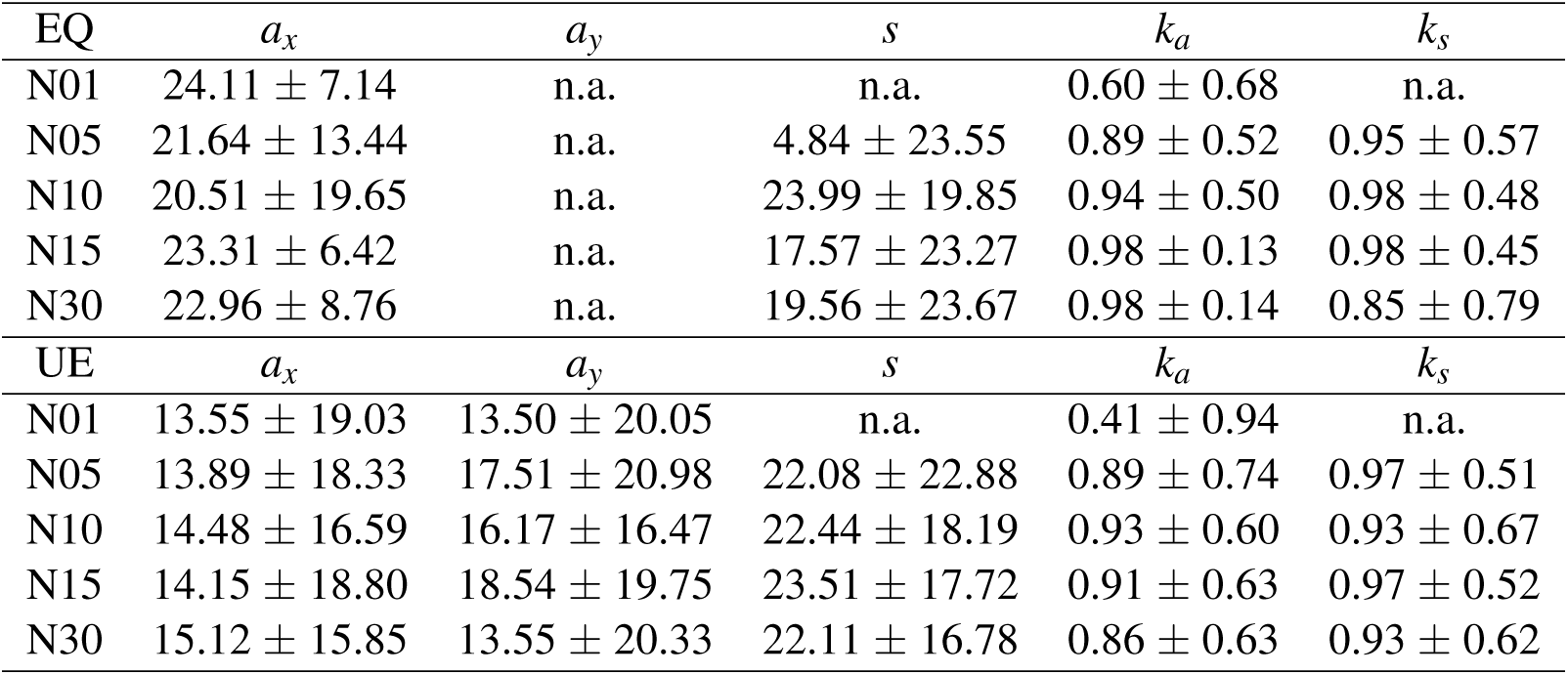
Averages (median±iqr) of best-fitting parameters per group size for prediction of food patch choice. Cases where parameters were not applied in the model are marked with n.a.

## Notes

### Competing Interest Statement

The authors have declared no competing interest.

